# Pattern similarity in the frontoparietal control network reflects an “off-veridical” template that optimizes target-match decisions during visual search

**DOI:** 10.1101/2021.12.18.473315

**Authors:** Xinger Yu, Joy J. Geng

**Affiliations:** Center for Mind and Brain, University of California, Davis, 267 Cousteau Pl., Davis, CA 95618 USA; Department of Psychology, University of California, Davis, One Shields Ave, Davis, CA 95616 USA

## Abstract

Theories of attention hypothesize the existence of an “attentional” or “target” template that contains task-relevant information in memory when searching for an object. The target template contributes to visual search by directing visual attention towards potential targets and serving as a decisional boundary for target identification. However, debate still exists regarding how template information is stored in the human brain. Here, we conducted a pattern-based fMRI study to assess how template information is encoded to optimize target-match decisions during visual search. To ensure that match decisions reflect visual search demands, we used a visual search paradigm in which all distractors were linearly separable but highly similar to the target and were known to shift the target representation away from the distractor features (Yu & Geng, 2019). In a separate match-to-sample probe task, we measured the target representation used for match decisions across two resting state networks that have long been hypothesized to maintain and control target information: the frontoparietal control network (FPCN) and the visual network (VisN). Our results showed that lateral prefrontal cortex in FPCN maintained the context-dependent “off-veridical” template; in contrast, VisN encoded a veridical copy of the target feature during match decisions. By using behavioral drift diffusion modeling, we verified that the decision criterion during visual search and the probe task relied on a common biased target template. Taken together, our results suggest that sensory-veridical information is transformed in lateral prefrontal cortex into an adaptive code of target-relevant information that optimizes decision processes during visual search.

## Introduction

When looking for a target object (e.g., a tiger hiding in the grasslands), we engage in a continuous “look-identify” cycle in which we use target features in memory (e.g., orange color) to guide attention and make match decisions. The memory representation of the target is referred to as the “attentional” or “target” template (Duncan & Humphreys, 1989), and its contents determine the efficiency of visual search (Geng & Witkowski, 2019; Hout & Goldinger, 2015; Malcolm & Henderson, 2009, 2010). The template is a core construct within all models of attention, but controversy remains over how target information is encoded in the brain. One possible reason for disagreement is that most target stimuli have simple static features, allowing for sensory and control regions to maintain veridical representations that are hard to distinguish from each other. However, recent behavioral studies have shown that under certain circumstances, target representations are shifted to be more optimal when “off-veridical” (Becker, 2010; Navalpakkam & Itti, 2007; Scolari et al., 2012; Yu & Geng, 2019), and these biased templates can be measured separately during attentional guidance and attentional decisions (Wolfe, 2021; Yu et al., in press; Yu & Geng, under review). In this study, we use such a paradigm to ask if biased target templates can be measured in frontoparietal and/or visual regions during a target decision task. We hypothesized that the need for a flexible, off-veridical target representation will result in greater reliance on lateral frontal regions that are known to dynamically code task-relevance (Duncan, 2001).

Lateral prefrontal cortex has long been hypothesized by attention researchers to hold the target template because neurons there selectively encode task-relevant information and maintain it over time (Funahashi et al., 1989; Fuster & Alexander, 1971; Miller et al., 1996). In one of the earliest demonstrations, Rainer et al., (1998) found that when shown an array of objects during a delayed-match-to-sample task, neurons in lateral prefrontal cortex maintained only the task relevant item over the subsequent delay period (Moore & Zirnsak, 2017; Squire et al., 2013). Since then, there have been many complementary studies in humans demonstrating that while task-relevant stimulus representations can be found widely throughout the brain (Christophel et al., 2017; Ester et al., 2015; Lee et al., 2013), the lateral prefrontal cortex specifically encodes the goal-relevant properties of the stimulus. For example, Long and Kuhl (2018) asked participants to make a judgment about a face stimulus based on a “goal cue” that indicated the dimension of relevance (gender or emotion). While decoding of stimulus properties in visual cortex was highly sensitive to sensory degradation following a visual mask, stimulus decoding within regions of the frontoparietal control network (FPCN; Schaefer et al., 2018) only occurred based on the goal-relevant stimulus dimension.

The data are consistent with the view that lateral prefrontal cortex encodes the contents of the task-based target template but guides attention and eye-movements through a functional network of oculomotor and sensory regions (Baldauf & Desimone, 2014; Bichot et al., 2015; Feredoes et al., 2011; Moore et al., 2003). Causal evidence for such network interactions was shown by Bichot et al. (2019) who found that silencing a region of ventrolateral prefrontal cortex impaired visual selectivity of visual search target features, suggesting this region maintains target templates that set attentional priority in sensory cortex. These studies are consistent with characterization of mid-lateral prefrontal cortex as being essential for control functions that selectively maintain information for visual search based on the current task demands (Badre & Nee, 2018; de la Vega et al., 2018; Duncan, 2001, 2013; Miller & D’Esposito, 2005; Panichello & Buschman, 2021).

Nevertheless, despite evidence implicating lateral prefrontal cortex in maintaining target representations, others have argued that low-level sensory regions may be better candidates for template storage because neurons in those regions are selectively tuned for different stimulus features (Harrison & Tong, 2009; Serences et al., 2009; Sreenivasan et al., 2014). However, many of these studies measured target representations during or just prior to target selection or comparison, leaving open the possibility that these measurements reflect “downstream” changes in sensory gain rather than the stable memory “source” of the target (Chelazzi et al., 1998; Desimone & Duncan, 1995; Reynolds & Heeger, 2009; Treisman & Gelade, 1980). Moreover, recent research has shown that information in target templates are not only used to modulate sensory processing but also used to determine target identity once an object has been selected (Bravo & Farid, 2016; Castelhano et al., 2008; Cunningham & Wolfe, 2014; Peltier & Becker, 2016; Rajsic & Woodman, 2020; Wolfe, 2021). This suggests that the target template contributes to attentional processes at multiple stages of processing and should be tuned to optimize target-to-distractor discriminability and minimize decision errors (Yu et al., in press).

To better understand how optimal template information is encoded, we conducted a visual search study with interleaved single-stimulus match-to-sample (probe) trials (Figure 1). The visual search task trained participants on the target feature relative to distractor features and the probe task asked participants to make target-match decisions. The separation of visual search and probe trials is essential for obtaining a measurement of target decisions without contamination from distractor processing and competition during active visual search given that target decisions more closely match the memory template (Yu et al., in press; Yu & Geng, under review). The task is based on previous work showing that difficult search for a target amongst linearly separable distractors (e.g., distractors are all “redder” than the orange target) induces a shift in the target template (e.g., becoming represented as yellower than the true target) (Bauer et al., 1996; Geng et al., 2017; Hodsoll & Humphreys, 2001; Scolari et al., 2012; Yu & Geng, 2019). Despite being off-veridical, the biased template is more optimal in this case because it prioritizes the ability to distinguish the target from distractors. The advantages in template-to-distractor distinctiveness have mostly been hypothesized to increase search efficiency by biasing sensory selection during attentional guidance (Becker, 2010; Navalpakkam & Itti, 2007; Scolari et al., 2012), but it has been recently shown in behavior to also facilitate target-match decisions (Yu et al., in press).

**Figure 1.**
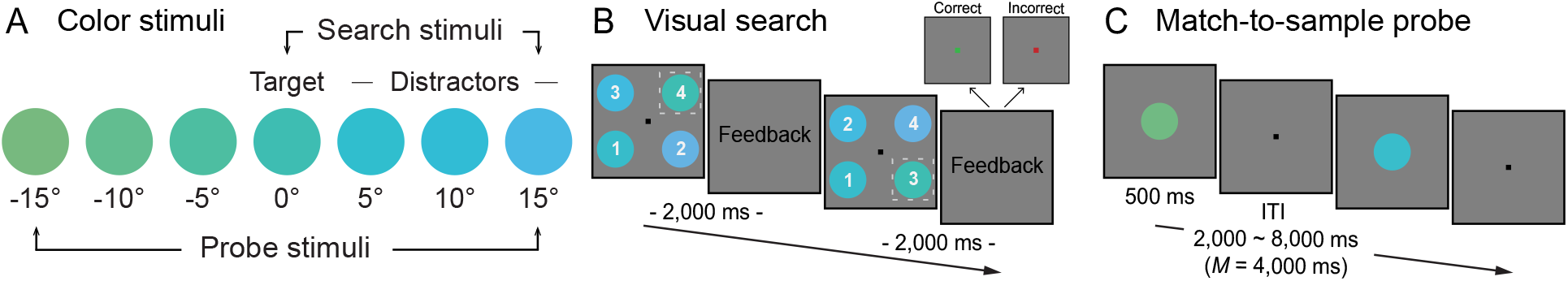
A) Color stimuli used in the fMRI experiment. The target color (0°) was a green-blue border color (Bae et al., 2015). The three distractors in visual search trials were all positive rotations (5°, 10° and 15°) from the target color. The probe colors included the target color (color 0°), and the three colors from each side of the target (i.e., ™5°, ™10°, ™15°, 5°, 10° and 15°). B) Example of visual search trials in the fMRI experiment. Participants were instructed to locate the target color circle and report the number within. Feedback was provided immediately after responses (green fixation squares for correct; red squares for incorrect). The white dashed squares illustrate the target but were not visible to the participants. C) Example of match-to-sample probe trials in the fMRI experiment. Participants were instructed to report if the centrally presented color circle was target color or not. There was no feedback in the probe trials. Non-target color values are exaggerated for visual clarity in all figures.

Here, we tested two non-mutually exclusive hypotheses concerning how the human brain encode the template information to optimize target-match decisions. Given the convergent findings from working memory, attentional control, and perceptual decision making, one hypothesis predicts that the off-veridical template representation would be stored and used to optimize target-match decisions in lateral prefrontal cortex within the frontoparietal network. Alternatively, a similar hypothesis could be made for visual cortex given evidence of working memory maintenance and target representations there during visual search (Chelazzi et al., 1998; Ester et al., 2009). We therefore identified a priori ROIs from frontoparietal control (FPCN) and visual (VisN) networks using a resting state network atlas defined from a large, independent sample of participants (Schaefer et al., 2018; Yeo et al., 2011). Evidence for a biased template even when there are no actual distractors present (on match-to-sample probe trials) would suggest that FPCN and VisN encode the target template as an adaptive memory of target characteristics in long-term memory to maximize the distinctiveness of target from distractors during visual search. Alternatively, if we find that the target representation is present, but veridical (i.e., centered around the true target value) on probe trials, it would suggest that the biased template is only invoked when it is actively used during visual search for attentional guidance. Furthermore, correspondence between pattern similarities in FPCN and VisN would suggest that a single template representation is used for target-match decisions; however, different patterns in FPCN and VisN would suggest that the readout of information from the sensory cortex is transformed in order to optimize the efficiency of target decisions.

## Results

### PFC encodes an off-veridical template to optimize target decisions

The goal of the study was to test whether the frontoparietal control and/or visual network encodes an off-veridical target template learned from optimizing target-match decisions during visual search. To do this, twenty participants performed a difficult *visual search* task and a match-to-sample *probe* task based on our previous work (Yu & Geng, 2019) while being scanned (Figure 1B). The search target was presented once at the beginning of the experiment as a colored item (e.g., a green-blue circle). All distractors were highly similar to the target (5°, 10°, and 15° from the target) but also linearly separable from the target (e.g., all being bluer than the target color based on unidirectional rotation on a continuous color wheel) (Figure 1A). Upon presentation of the display, participants searched for the predefined target-color circle and reported the number inside by pressing 1-4 on a button box. The visual search trials were used to train participants’ expectations for the target with respect to distractor colors (Becker, 2010; Navalpakkam & Itti, 2007; Yu & Geng, 2019). Overall performance was high (accuracy: *mean* ± 95% CI = 88% ± 4%; RT: *mean* ± 95% CI = 812ms ± 85ms) suggesting that participants had a target template that could be successfully distinguished from distractors (Supplemental Figure 1). The findings replicate our previous results using a similar paradigm (cf. data from Experiment 1, Yu & Geng, 2019).

On separate match-to-sample *probe* trials (Figure 1C), participants indicated whether or not a particular color stimulus (0°, ±5°, ±10° and ±15° from the target) (Figure 1A) was the target (Geng et al., 2017; Yu & Geng, 2019). Because each probe trial had only one stimulus presented at the center of the screen (Figure 1C), we were able to obtain a pure measurement of match decisions independent from rapid visual search cycles of target localization and identity decisions. Previous behavioral work has demonstrated that under these conditions, the target representation shifts “off-veridical” away from distractors such that targets with the relational (e.g., “greener”) values (i.e., negative probe colors) were more likely to be selected and moreover, this bias was preserved stably in memory (Yu et al., in press). The data of primary interest for the fMRI analyses therefore included only the probe trials and are described in detail below.

#### Behavioral representational geometry

The mean proportion of “target yes” responses for each of the seven probe stimuli are shown in Figure 2A. The false alarm rates of non-target colors, as a measurement of which non-target colors were mistaken as the target color, were entered into a paired sample *t* test to test for differences between color directions (negative, positive). Participants had significantly more false alarms to the negative probe colors (*mean* ± 95% CI = 44% ± 10%) compared to the positive ones (*mean* ± 95% CI = 12% ± 7%), t_19_ = 4.55, *p* = .0002, *d* = 1.02, BF_10_ = 139.58. The behavioural results replicated previous findings (Navalpakkam & Itti, 2007; Scolari et al., 2012; Yu & Geng, 2019) that the target representation shifted away from distractors to enhance optimal off-target features that increase the template-to-distractor distinctiveness (Geng & Witkowski, 2019).

**Figure 2.**
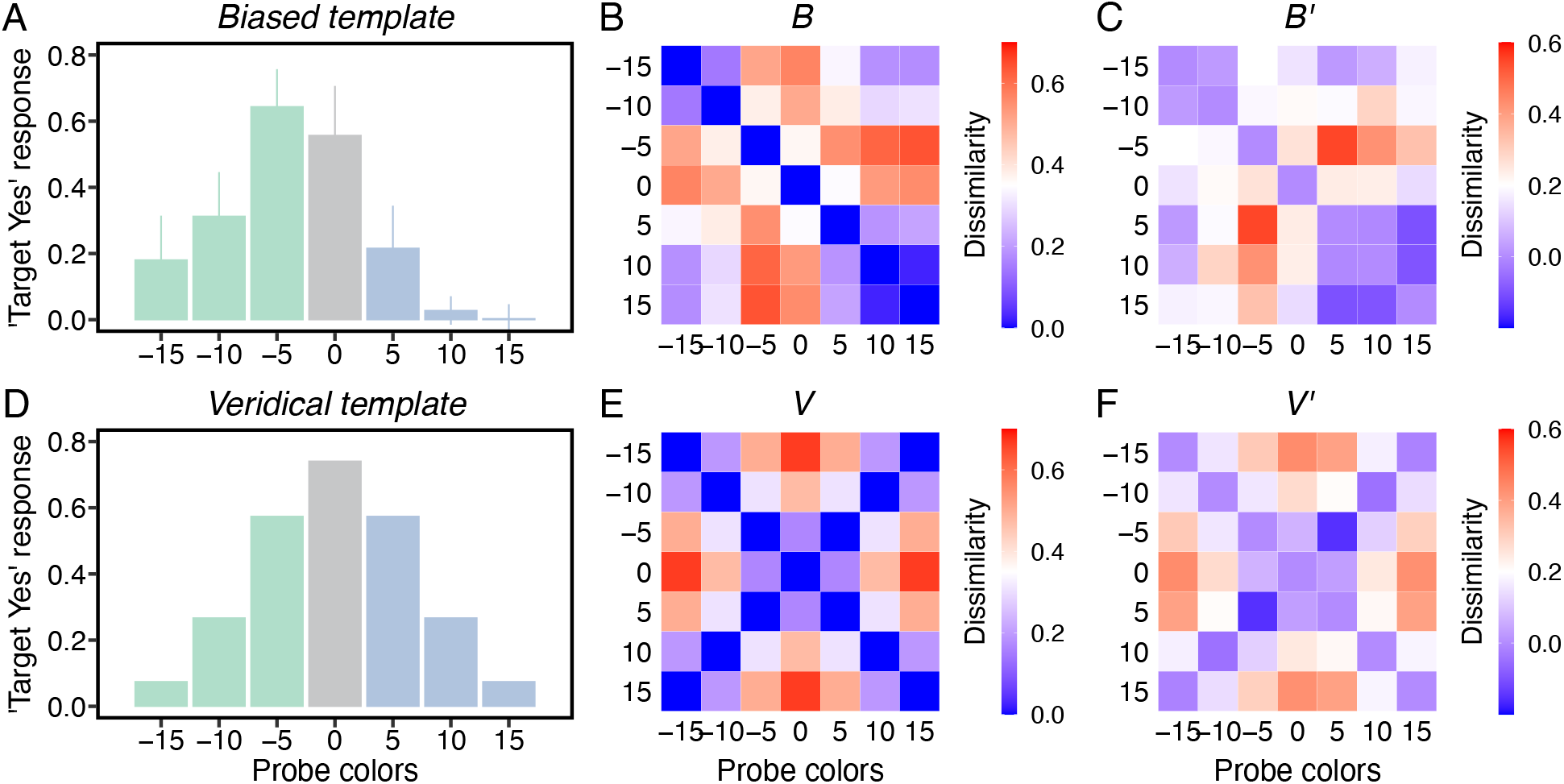
A) Group average of “target yes” responses. The center gray bar indicates proportion of “hits” in response to the true target color and other colored bars indicate “false alarms” to non-target colors. All error bars are 95% confidence intervals. B) The biased behavioral RDM averaged across participants (*B*). This RDM was generated by participants’ own probe task performance and reflected the biased target representation in Figure A. See Supplemental Figure 2 for individual RDMs. C) To ensure the effects were specific to each of the model RDMs (*B* and *V*), we conducted a partial correlation analysis to regress out their covariance with the other. The partial biased matrix (*B’*) highly correlates with *B* but not with *V*. D) The veridical target template. This template is a theoretical template, which assumes the target template was normally distributed and centered at the target color. E) The veridical behavioral RDM (*V*). F) The partial base matrix (*V’*) has a strong correlation with *V* but extremely weak correlation with *B*. Redder colors indicate more dissimilarity between two stimulus conditions (e.g., -5° and 15°); bluer colors indicate more similarity between two stimulus conditions (e.g., 10° and 15°).

Next, we quantified the representational distance between each pair of probe stimuli by constructing a representational dissimilarity matrix (RDM) for each participant from their own data. This matrix, denoted (*B*), reflected each individual’s biased target template (see Figure 2B for group average RDM and Figure S2 in Supplemental Material for individual RDMs) and was created as a model for subsequent analyses of brain data. In addition to the biased RDM (*B*), we built a veridical RDM (*V*) (Figure 2E) to serve as an alternative baseline model of an unbiased target template (Figure 2D). To ensure that *B* and *V* capture distinct patterns, we performed a partial correlation analysis to regress out their shared covariance (Park et al., 2020). As expected, the partial biased model RDM (*B’*) did not correlate with *V* (*mean* ± 95% CI = -.03 ± .03, z_19_ = - 2.28, *p* = .99), but did correlate strongly with *B* (*mean* ± 95% CI = .55 ± .12, z_19_ = 3.92, *p* < .0001) (Figure 2C). In contrast, the partial base model RDM (*V’*) had a high correlation with *V* (*mean* ± 95% CI = .64 ± .08, z_19_ = 3.92, *p* < .0001) but only a weak one with *B* (*mean* ± 95% CI = .06 ± .03, z_19_ = 2.91, *p* = .001) (Figure 2F).

#### Brain representational geometry

To test our primary hypotheses, we first test for representations of the target in our *a priori* anatomical ROIs based on FPCN and VisN networks (Figure S3 in Supplemental Material) defined through resting state data (Figure 3A): bilateral prefrontal-FPCN (PFC), parietal-FPCN (PAR) and central-VisN (VIS). The representational distances (one-sided Wilcoxon signed rank test; Holm-Bonferroni corrected from multiple comparisons across numbers of model RDMs, *N* = 2, and bilateral ROIs, *N* = 8) estimated in bilateral PFC were explained by the model RDM *B’* (left: *mean* ± 95% CI = .12 ± .07, z_19_ = 3.10, *p* = .007; right: *mean* ± 95% CI = .10 ± .06, z_19_ = 3.02, *p* = .009), but not by *V’* (left: *mean* ± 95% CI = .02 ± .06, z_19_ = .56, *p* = .89; right: *mean* ± 95% CI = .03 ± .06, z_19_ = 1.16, *p* = .52). In addition, the Kendall’s τ_A_ rank correlations between the *B’* and *V’* model RDM and the brain RDM in bilateral PFC were compared using a one-sided paired samples Wilcoxon test to determine if the correlation with *B’* and *V’* differed from each other statistically. Significant differences were found in left PFC (z_19_ = 1.72, *p* = .044), but not in right PFC (z_19_ = 1.12, *p* = .14). Notably, while left PAR was significantly correlated with *B’* (*mean* ± 95% CI = .09 ± .06, z_19_ = 2.76, *p* = .025), the right PAR pattern was only explained by *B’* at a reduced threshold (*mean* ± 95% CI = .08 ± .07, z_19_ = 2.17, *p* = .015, uncorrected) (Figure 3B). The representations in bilateral PAR were not explained by *V’* (left: *mean* ± 95% CI = .04 ± .05, z_19_ = 1.23, *p* = .58; right: *mean* ± 95% CI = .04 ± .06, z_19_ = 1.32, *p* = .61). The Kendall’s τ_A_ rank correlations with *B’* were not significantly higher than the correlations with *V’* in bilateral PAR (left: z_19_ = .78, *p* = .23; right: z_19_ = .71, *p* = .25). In contrast, the representations in bilateral VIS were explained by *V’* (left: *mean* ± 95% CI = .10 ± .04, z_19_ = 3.58, *p* = .001; right: *mean* ± 95% CI = .10 ± .05, z_19_ = 3.21, *p* = .004), but not by *B’* (left: *mean* ± 95% CI = .03 ± .04, z_19_ = 1.46, *p* = .62; right: *mean* ± 95% CI = .04 ± .05, z_19_ = 1.46, *p* = .54) (Figure 3B). The Kendall’s τ_A_ rank correlations with *V’* were significantly higher than the correlations with *B’* in right VIS (z_19_ = 1.68, *p* = .048), but only marginally significantly in left PAR (z_19_ = 1.46, *p* = .077). To be sure that these findings were specifically related to our task, we further looked for brain-model pattern similarity within a control region in bilateral auditory cortex (AUD) defined by the same Schaefer atlas (Schaefer et al., 2018; Yeo et al., 2011). The pattern similarity in bilateral AUD was not explained by either template model RDM (z_19_ < 2.05, *ps* > .2) (Figure 3B).

**Figure 3.**
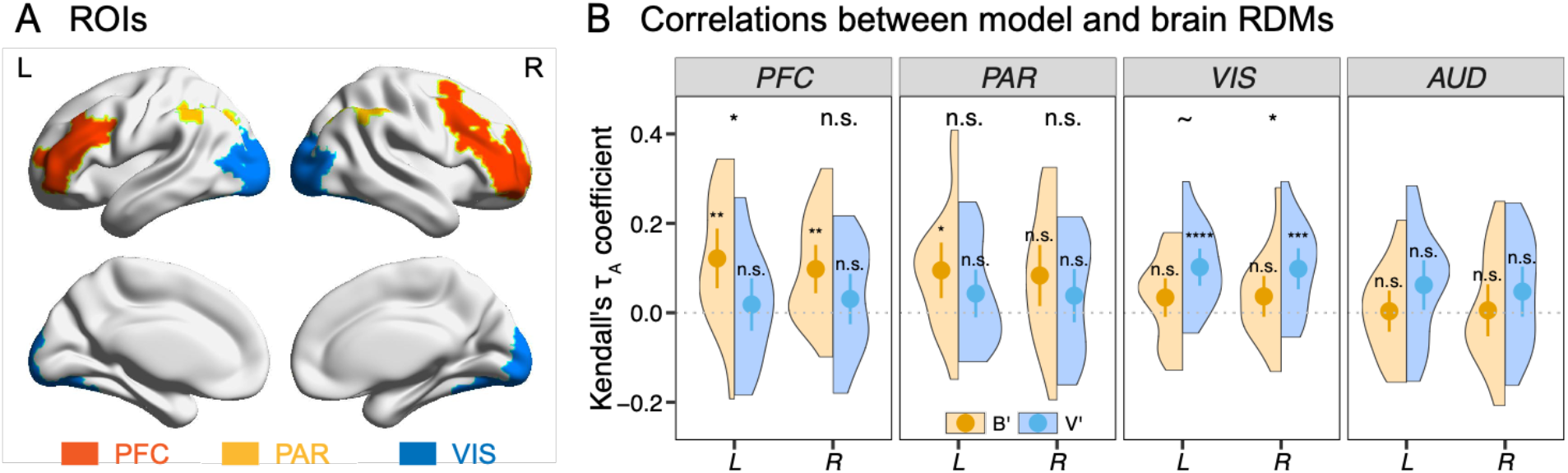
A) We assessed target representations in two resting-state networks defined from a large, independent sample of participants (Yeo et al., 2011): the frontoparietal control network (FPCN) and the visual network (VisN). FPCN was divided into discontinuous prefrontal and parietal subregions, and VisN was separated into central (i.e., striate and extrastriate cortex) and peripheral subregions. All the cortical parcels corresponding to the prefrontal-FPCN (PFC), the parietal-FPCN (PAR) and the central-VisN (VIS) were selected as ROIs. B) Mean Kendall’s τ_A_ rank correlations between the model RDMs (*B’* and *V’*) and the brain RDM. The significance of the correlations was assessed by the one-sided Wilcoxon signed rank test. \*\*\*\**p* < .001 (Holm-Bonferroni corrected from multiple comparisons across numbers of model RDMs, *N* = 2, and bilateral ROIs, *N* = 8), \*\*\**p* < .005, \*\**p* < .01, \**p* < .05, ^∼^*p* < .1, and n.s. > .1. All error bars are 95% confidence intervals.

The differences in PFC and VIS illustrate a “double dissociation” in which the prefrontal cortex matched the biased model better than the veridical one and the opposite pattern was found in the early visual cortex. The fact that the visual ROI encoded the veridical target, but the prefrontal cortex encoded the biased template during the same probe trial suggests that match decisions transformed veridical visual representations into a template space designed to improve target distinctiveness from visual search distractors. Furthermore, that this transform occurred on the probe trials when distractors were absent suggests that target decisions are based on a stable biased target representation stored in long-term memory (Wolfe, 2021; Yu et al., in press; Yu & Geng, under review).

#### Whole brain searchlight analysis

In order to capture all of the brain regions that possibly encode the target template, we conducted a whole brain searchlight analysis for each participant using their own biased RDM (*B*) (Figure S2 in Supplemental Material) and the baseline veridical RDM (*V*) (Figure 2E) as separate models. The group analysis with *B* identified five significant brain regions located in bilateral middle frontal gyrus (MFG) (left: [x, y, z] = [-32, 40, 14], z_19_ = 5.43; right: [x, y, z] = [34, 42, 32], z_19_ = 4.66), left superior parietal lobe (SPL) ([x, y, z] = [-38, -42, 32], z_19_ = 4.58), left percental gyrus (PrG) ([x, y, z] = [-46, -14, 48], z_19_ = 5.92), and right supplemental motor cortex (SMC) ([x, y, z] = [8, 20, 44], z_19_ = 4.86) (Figure 4A). The significant bilateral MFG result corresponds to the PFC ROI in FPCN from the previous hypothesis driven analysis. The group analysis with *V* also identified five significant brain regions located in bilateral occipital gyrus (OcG) (left: [x, y, z] = [-16, -96, 8], z_19_ = 5.13; right: [x, y, z] = [20, -90, 18], z_19_ = 4.95), replicating the hypothesis driven results above, and also left precentral gyrus (PrG) ([x, y, z] = [-46, 2, 36], z_19_ = 4.76), left postcentral gyrus (PoG) ([x, y, z] = [-40, -18, 50], z_19_ = 4.85), and left supplemental motor cortex ([x, y, z] = [-14, -2, 62], z_19_ = 4.63) (Figure 4B). Statistics and the MNI coordinates of the peak voxels in the five clusters are reported in Table 1. The significant results observed in motor regions, e.g., left PrG (contralateral to the right index and middle fingers that were used for button presses), were due to the actual motor response rather than response independent decision processes (see Supplemental Material button response RDM searchlight).

**Figure 4.**
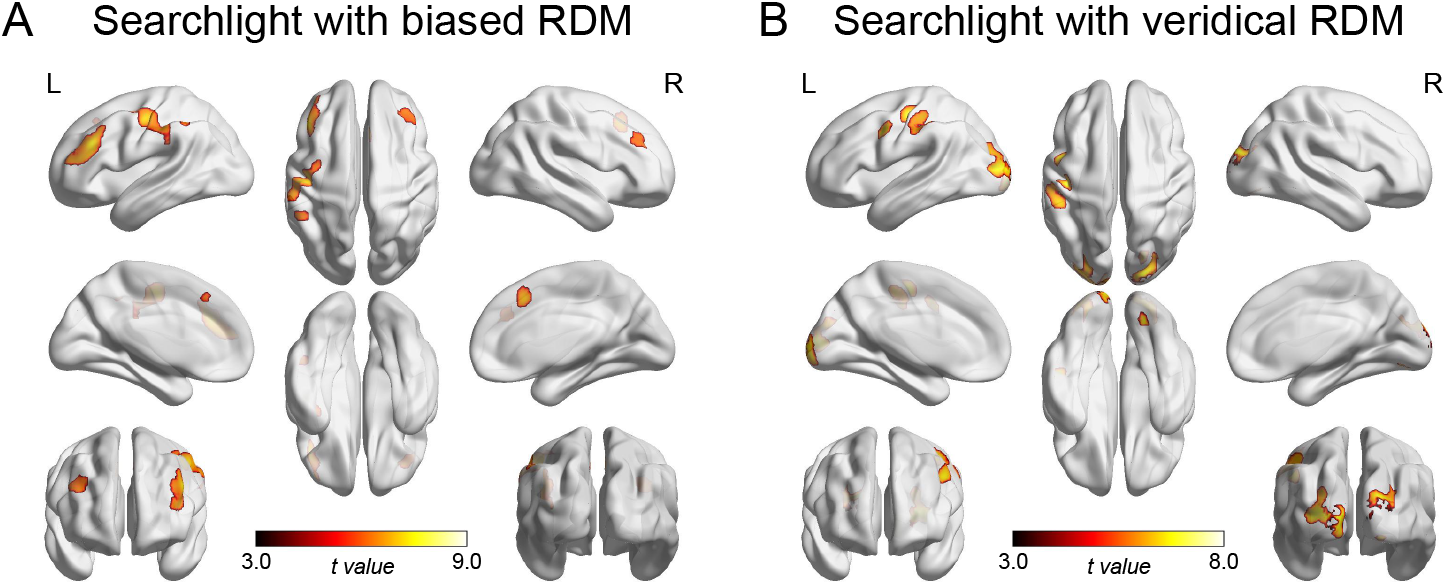
A) Brain maps showing the clusters identified in the whole brain searchlight analysis with the biased RDM (*B*). B) Brain maps showing the clusters identified in the whole brain searchlight analysis with the veridical RDM (*V*).

**Table 1.** Brain regions in which the neural activity pattern similarity was significantly correlated with the behavioral RDMs in the searchlight analysis.

| Biased behavioral RDM | Veridical behavioral RDM |
| --- | --- |

| Anatomical regions | x | y | z | # voxels | z value | Anatomical regions | x | y | z | # voxels | z value |
| --- | --- | --- | --- | --- | --- | --- | --- | --- | --- | --- | --- |
| L MFG | -32 | 40 | 14 | 1353 | 5.43 | L OcG | -16 | -96 | 8 | 1155 | 5.13 |
| R MFG | 34 | 42 | 32 | 364 | 4.66 | R OcG | 20 | -90 | 18 | 1180 | 4.95 |
| L SPL | -38 | -42 | 32 | 184 | 4.58 | L PoG | -40 | -18 | 50 | 852 | 4.85 |
| L PrG | -46 | -14 | 48 | 951 | 5.92 | L PrG | -46 | 2 | 36 | 287 | 4.76 |
| R SMC | 8 | 20 | 44 | 382 | 4.86 | L SMC | -14 | -2 | 62 | 186 | 4.63 |

Coordinates (x, y and z) are reported in MNI space

### The biased decision template encoded in PFC was used during active visual search

We concluded from the fMRI results that visual cortex contained a veridical copy of the target template, but lateral prefrontal cortex encoded target decisions based on an off-veridical template in long-term memory biased by the search context. However, because the fMRI target identification task relied solely on probe trials, it is possible that the biased decision template we measured in PFC is different from that used during active visual search. Although it is not possible to isolate target-match decisions during visual search using fMRI given the rapid cycles of object selection and target-match evaluations before the target is found (Yu et al., in press), we tested for correspondence in decision processes on visual search and probe trials using drift diffusion modeling (DDM). We conducted an online behavioral study in which we applied DDM to characterize how accurately and quickly the target was selected from distractors during active visual search (Figure 5B, 5D) and how accurately and quickly the probe stimulus was identified as a target or non-target on probe trials (Figure 5C, 5E). Each visual search trial consisted of two bilaterally presented target and distractor circles (Figure 5A). *Standard* visual search trials always contained a positively 10° rotated distractor color and were used to set up expectations regarding the distractor context. On *critical* search trials, the distractors varied from the target color from +/-30° in steps of 5°. *Critical* distractors, unlike *standard* distractors, could be positive or negative rotations from the target color and provided a way to assess how well each color matched the internal target template used for decision making during active visual search. The *probe* trials were identical to the fMRI experiment and asked participants to indicate whether a particular color stimulus (ranging from -30° – +30°) was the target (Figure 5A). A high correlation between individual drift rates on the *critical* search trials and drift rates from the *probe* trials would suggest that the two processes rely on a similar underlying decision criterion based on the target template.

**Figure 5.**
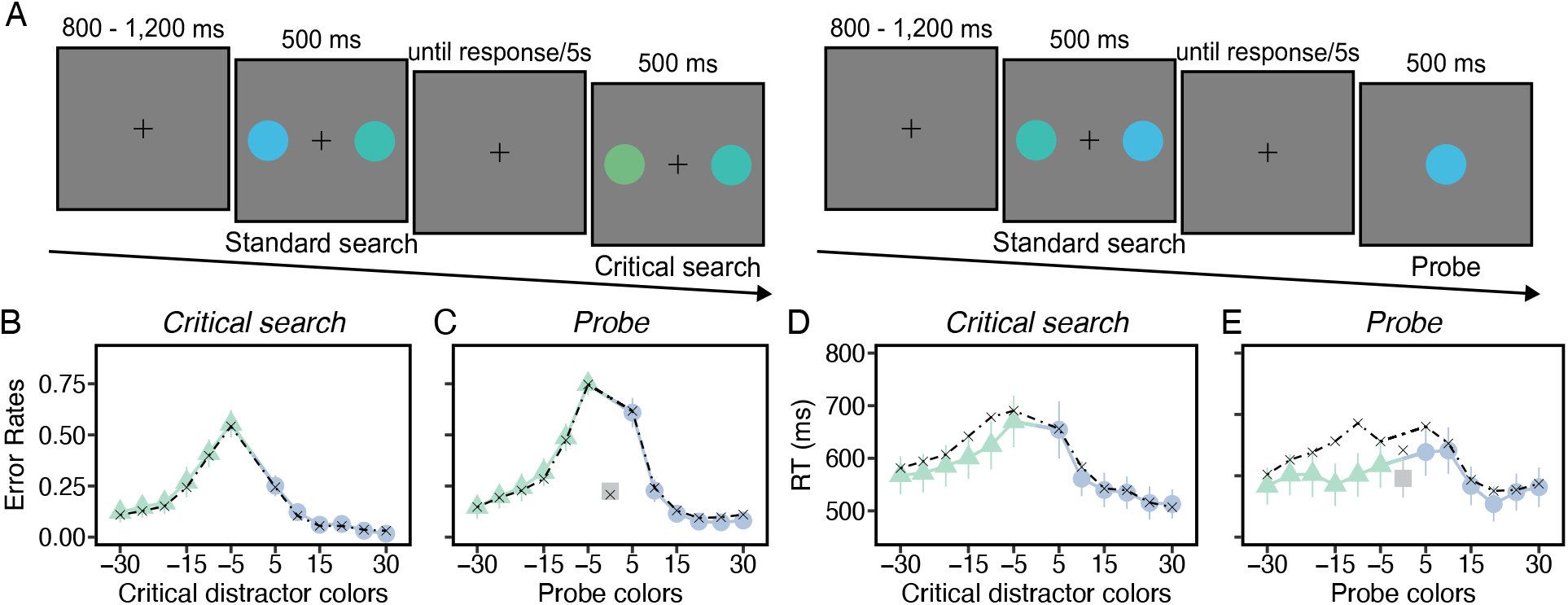
A) Example of *standard* and *critical* visual search trials, and *probe* trials. On search trials, participants were instructed to locate the target color circle and press a mouse button indicating its location. On probe trials, participants were instructed to report if the centrally presented color circle was the target color or not. No feedback was given on any of the trials. B) Error rates from the critical visual search trials. C) Error rates from the probe trials. D) RTs from the critical search trials. E) RTs from the probe trials. Green and blue hues indicate negative and positive values, respectively, on the x-axis (gray indicates 0°). The black dashed lines represent the “fitted curves” for error rates and RT from the DDM for the best fit DDM parameters. All error bars are 95% confidence intervals.

#### Analysis of the drift rates

Figure 6A and 6B show the group mean posterior estimates of the drift rates for the visual search and probe conditions, respectively. If the probe task taps into target information that is used during visual search decisions, we expected a positive correlation between these two metrics. Following the analysis strategy from the fMRI experiment, each participant’s drift rates were converted to a dissimilarity matrix (Figure 6C and 6D). The significance of correlations between the search drift rate RDM and the probe drift rate RDM was evaluated by a permutation test by randomizing stimulus labels. There was a significantly positive correlation between the two RDMs, *mean* ± 95% CI = .55 ± .04, *p* = .0004, highlighting the close relationship between the decision processes engaged during probe trial judgements of target color held in memory over time, and the decision processes engaged during active comparisons of the target template to search items during visual search.

**Figure 6.**
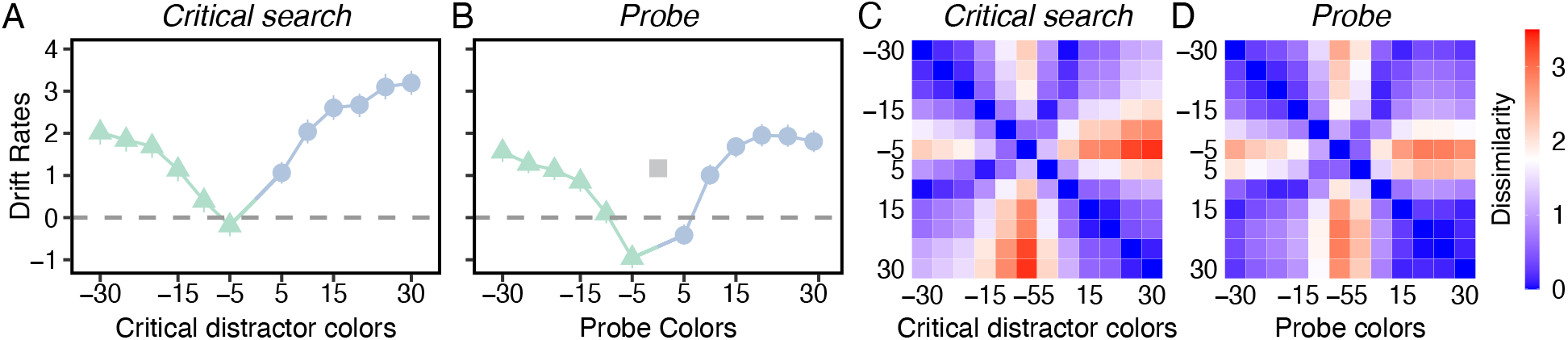
A) The drift rates for different critical search conditions. B) The drift rates for different probe color conditions. C) The search drift rate RDM averaged across participants. D) The probe drift rate RDM averaged across participants. All the error bars are the 95% confidence intervals.

Importantly, both drift rates showed the same biased pattern we observed in the fMRI experiment in FPCN. In particular, as shown in Figure 6A and 6B, the drift rate was negative at the -5° distractor in both critical search and probe trials; this indicates that the accumulation of evidence was slow, and moreover, that the “wrong” decision that the -5° stimulus was the target, was made more often than not on both visual search and probe trials. This pattern is consistent with the use of an off-veridical template to make target decisions during both visual search and probe trials. Also consistent with the notion that the biased template was created to increase target-to-distractor distinctiveness during visual search, the drift rates show that the “target no” response decision times were faster for the positive colors (*mean* ± 95% CI = 551ms ± 29ms) that were seen as actual visual search distractors compared to the negative colors (*mean* ± 95% CI = 602ms ± 36ms) (t_69_ = -7.05, *p* < .0001, *d* = -.84, BF_10_ > 1,000) (Figure 5D). These results support those from the fMRI experiment and suggest that the decision processes during both visual search and probe trials relied upon a common biased target template in long-term memory that reflects a mechanism to increase the psychological distance between the target and distractors (Geng & Witkowski, 2019; Yu et al., in press).

## Discussion

Models of attention theorize the existence of an internal representation of target features in memory that is referred to as an attentional or target template (Bundesen, 1990; Duncan & Humphreys, 1989; Wolfe, 2021). It is commonly believed that lateral prefrontal cortex maintains template information to direct attention and eye-movements to possible targets through a functionally connected network of oculomotor and sensory regions (Baldauf & Desimone, 2014; Bichot et al., 2015; Feredoes et al., 2011; Moore et al., 2003). However, less is known about how template information might be used to optimize target-match decisions once a potential target has been selected. Here, using pattern-based fMRI methods and a perceptual decision-making task, we compared representations of target features across two resting-state networks. We targeted networks for which there is evidence of template maintenance, namely the frontoparietal control network and the visual cortical network (Schaefer et al., 2018; Yeo et al., 2011).

The argument for the importance of decision processes during visual search is based on the “look-identify” cycle inherent in visual search: Once a candidate object has been localized by attention, a decision must be made regarding the exact identity of the stimulus as a match or non-match to the target (Castelhano et al., 2008; Hout & Goldinger, 2015; Rajsic & Woodman, 2020). This decision component is a time-consuming part of the visual search cycle and must be accurate if visual search is to be ultimately successful. We previously showed that target-match decisions operated on more precise template information than attentional guidance, presumably because decisions need to be highly accurate whereas initial selection can be less precise (Yu, et al., in press; Yu & Geng, under review). Similarly, Rajsic & Woodman (2020) found that the benefit of weighting the target template over accessory memory representations in a dual-memory search task mainly lies in enhancing target recognition rather than more efficient localization of the target, again presumably because recognition serves to “gate keep” the outcome accuracy of object localization. Moreover, template information used during the decision stage more closely matched the precision of target information in long-term memory than guidance (Yu, et al., in press; Yu & Geng, under review). Together, these suggest that attentional decisions are better measurements of the underlying precision of the target template in memory than guidance.

In our current study, we successfully decoded the off-veridical template that maximized the target-to-distractor discrimination in PFC during a target identification probe task that isolated the match decisions from other processes of visual search. The results suggest that PFC engages in target decisions using templates that are held in long-term memory and shaped by the recent search context. This is consistent with previous research that frontoparietal regions dynamically change target representations depending on task contexts (Bracci et al., 2017; Ester et al., 2016; Lee et al., 2013; Long & Kuhl, 2018; Sarma et al., 2016; Swaminathan & Freedman, 2012). This information in lateral prefrontal cortex is then used to make target-match decisions, in addition to setting sensory priority to guide attention.

In contrast to the pattern in FPCN, our data indicates that stimulus responses in early visual cortex reflected the veridical target. The correlations between each brain region’s responses and the two target RDMs (biased, veridical) illustrate a double dissociation that suggests that match decisions were based on a transformation of veridical target information in sensory cortex into a biased template space in service of improving decisions about target identity. This transformation suggests that improved visual search performance can be achieved by flexibly changing sensory readout, while leaving the sensory representations unchanged (Birman & Gardner, 2019).

In the online behavioral study, we found a close relationship between the target-match decisions in visual search and probe trials by applying DDM to accuracy and RT in the two tasks. This suggests that the decision process engaged during active visual search and the decision process engaged during probe trial judgments of target color rely on a similar underlying accumulation of evidence based on an off-veridical template. The fact that the biased template was used to make decisions on probe trials when no distractors were actually present, indicates that experience with the target within a linearly separable context produces a change in the memory representation of the target used for making decisions about identity (Yu & Geng, 2019). This is consistent with the preferential decoding of the biased template in PFC, given that lateral prefrontal cortex adaptively encodes task relevant information in working memory (D’Esposito & Postle, 2015; Duncan, 2001; Lee et al., 2013; Miller et al., 1996; Riggall & Postle, 2012).

In conclusion, our findings provide evidence that the frontoparietal control network represents the target template as an adaptative memory of target features to optimally discriminate the target from distractors; in contrast, early visual cortex encodes veridical inputs, suggesting that decision processes about target identity are based on a transformation of veridical information when doing so increases target discrimination. This work strongly supports the role of lateral prefrontal cortex in flexibly controlling target relevant information in accordance with task goals to optimize decision processes during visual search.

## Materials and methods

### Participants

#### fMRI experiment

Twenty-one participants (self-reported 8 males and 13 females, all right-handed, aged between 19-30 years) participated in a 1.5h scanning session and received monetary compensation. Data from one participant were excluded from analyses due to excessive head motion larger than 4mm^3^, which resulted in a final group of twenty participants (self-reported 7 males, 13 females). This number far exceeded the estimated sample size of *N* = 7 (.95 power, .05 two-tailed significance) calculated with G*power (http://www.gpower.hhu.de/) based on behavioral data from our previous experiment that used similar stimuli and procedures (Yu & Geng, 2019; Experiment 1; unidirectional group). The behavioral effect we wished to detect was highly reliable, but we set our sample size at 20 due to expected noise associated with fMRI measurements. Each participant provided written informed consent in accordance with the local ethics clearance as approved by the National Institutes of Health. Each participant’s color vision was assessed by self-report on an online color blindness questionnaire (https://colormax.org/color-blind-test). All participants had normal or corrected-to-normal vision and no history of neurological or psychiatric illness.

#### Online experiment

Seventy participants (self-reported 10 males and 60 females, 9 left-handed and 61 right-handed, aged 18-30 years) from University of California, Davis participated online in partial fulfillment of a course requirement. The sample size *N* = 70 (.85 power, .05 two-tailed significance) was determined based on a previous online study from which we adopted the experimental design (Yu, et al., in press; online Experiment 2). Eight participants were excluded from data analyses due to poor visual search performance (i.e., accuracy in standard visual search trials was below 75%). Data were collected until we obtained the target sample of 70 participants after exclusion criteria were applied. Each participant provided written informed consent in accordance with the local ethics clearance as approved by the National Institutes of Health. Each participant’s color vision was assessed by self-report. All participants had normal or corrected-to-normal vision, and all had normal color vision.

### Task and Stimuli

#### fMRI experiment

All stimuli were generated by a Dell computer and displayed on a 24” BOLDscreen LCD monitor with a spatial resolution of 1920 × 1200 pixels. The operating system was Windows 7, and Matlab Psychtoolbox (Brainard, 1997; Pelli, 1997) was used to create stimuli. The target and distractor colors were selected from a color wheel defined in CIELAB color space (a, b coordinates = 0, 0; luminance = 70). We used a green-blue boarder color (190°) and a blue-purple border color (274°) as the target colors to control for the potential color category effects on responses (Bae et al., 2015). The two target colors were counterbalanced across participants. Each participant was assigned a single target color throughout the experiment. Although the two target colors appeared to affect visual search RT differently (t_19_ = -4.20, *p* < .001), they did not impact accuracy (*p* = .29) nor performance on the probe task (*p* = .72). Data from these conditions were therefore collapsed to maximize power. The three distractors in each search trial were different from each other and always 5°, 10° and 15° positively rotated away from the target color. These distractors were chosen to exceed the average just noticeable difference, but still be confusable with the target when presented in a competitive search context (Geng et al., 2017; Yu & Geng, 2019). The colors in probe trials included the target color (0°), and three non-target colors from each side of the target (i.e., ±5°, ±10° and ±15°).

An example of the target color was presented at the beginning of the experiment. On visual search trials (Figure 1C), four circles (3° of visual angle in diameter) were presented for 1000ms on a gray background. The target color was always present and was randomly located at one of the four vertices along an imaginary square (6° of visual angle from center to edge); distractors appeared at the other three vertices. A number from 1-4 (1° of visual angle; white) was centrally located within each circle without duplication. Upon presentation of the search display, participants located the pre-defined target color circle and reported the number inside by pressing 1-4 on a button box with their right hand. Visual feedback was provided immediately following responses (the fixation square turned green for correct responses and red for incorrect responses). Each match-to-sample *probe* trial consisted of a centrally presented circle (3° of visual angle in diameter) for 500ms (Figure 1C). Participants reported whether the circle color was the target color: “yes” responses were indicated by a button press with the right index finger and “no” responses were indicated by a button press with the middle finger. Participants were informed that no feedback was provided on probe trials because there was no absolute “correct” or “incorrect” answer. If no response was made within 2000ms, both visual search and probe trials automatically terminated.

Prior to the start of the experiment, participants completed 48 practice trials composed of both search and probe trials. Participants were instructed to fixate on the center fixation square (.17° of visual angle from center to edge) throughout the experiment. The main experiment was composed of 320 visual search trials and 448 probe trials. Trials were presented in 8 blocks, each containing 2 alternating blocks of visual search trials and probe trials. Each alternating block contained 20 visual search trials followed by 28 probe trials. There was a 10s inter-trial interval when switching between visual search trials and probe trials. The probe trials in each scan had 24 repetitions of 0°, 16 repetitions of ±5°, 8 repetitions of ±10°, and 8 repetitions of ±15°. We used an uneven ratio (1:2) of veridical targets to non-target foils in order to maximize the number of non-target color presentations and to roughly equate the number of likely “yes” and “no” responses to avoid response biases, based on our previous findings (Yu & Geng, 2019).

#### Online experiment

The experiment was conducted entirely online through Testable (https://www.testable.org/). All stimuli were created in Illustrator, saved as PNG files, and uploaded to Testable.org. All stimuli were presented against a gray background (color hue = ‘#808080’). The two target colors (190° and 274°) were identical to the fMRI experiment and counterbalanced across participants. Because there were no spurious differences (*ps* > .35), the data were collapsed to maximize power in all subsequent analyses. There were three types of trials: 1) *standard* visual search trials to set up expectations for the distractor colors; 2) *critical* search trials to assess how target templates are used to identify targets during visual search; 3) *probe* trials to measure the target template independent of simultaneous distractor competition.

Each visual search trial (Figure 5A) consisted of two bilateral target and distractor circles (radius: 135 pixels) on the horizontal meridian (distance between the centers of the two circles: 400 pixels). The distractor color in the *standard* visual search trials was always positively 10° rotated from the target color. The *critical* distractor set was constructed in steps of 5° from the target color to +/-30° rotations from the target color, resulting in a total of 12 critical distractor colors. Each *probe* trial consisted of a centrally presented circle (radius: 135 pixels) (Figure 5A). The colors in probe trials included the target color (0°), and twelve non-target colors identical to the critical distractor set.

An example of the target color was presented at the beginning of the experiment. On search trials, participants were instructed to indicate whether the target color appeared on the left side by pressing the left arrow key or on the right side by pressing the right arrow key. On probe trials, participants were required to report whether the circle color was the target color by pressing the left arrow key for “yes” responses or the right arrow key for “no” responses. The stimuli in both trials appeared on the screen for 500ms and participants had up to 5s to make their responses. No feedback was given. After response, a central fixation cross was presented for 800-1200ms before the next trial started.

Participants completed 20 practice standard visual search trials with feedback before the main experiment started. Participants were instructed to fixate on the center cross when no stimuli were presented on the screen. The main experiment was composed of 108 standard visual search trials, 72 critical search trials and 108 probe trials. As in the fMRI experiment, we used an uneven ratio (1:2) of veridical targets to non-target foils to maximize the number of non-target color presentations and to roughly equate the number of likely “yes” and “no” responses. Trials were presented in 12 blocks. All blocks started with 9 standard visual search trials, after which 12 critical search trials presented in half of the blocks and 18 probe trials presented in the other half.

### Regions of interest selection

We assessed target representations in two resting-state networks defined from a large, independent sample of participants (Yeo et al., 2011): the frontoparietal control network (FPCN) and the visual network (VisN) (Figure S3 in Supplemental Material). The 400 brain parcels (Schaefer et al., 2018) were projected onto the high-resolution anatomical image of each participant using the FreeSurfer cortical parcellation scheme (http://surfer.nmr.mgh.harvard.edu). Each parcel was matched to one of the seven functional networks identified by Yeo et al., 2011. Next, we decomposed FPCN into separate prefrontal and parietal subregions and divided VisN into central (i.e., striate and extrastriate cortex) and peripheral subregions. All the cortical parcels corresponding to the prefrontal-FPCN (PFC), the parietal-FPCN (PAR), and the central-VisN (VIS) were selected as regions of interest (ROIs) for the representational similarity analysis (Figure 3A). The mean number of voxels in each ROI are reported in Table 2. None of the voxels from one ROI were contiguous with voxels from another ROIs. We also included the bilateral auditory cortex (AUD) in the somatomotor network as control regions. All ROIs were then coregistered to the functional data.

**Table 2.** Mean number of voxels in each region of interest selected from the Schaefer atlas.

| Region of interest | Hemisphere | # of voxels |
| --- | --- | --- |
| PFC | L | 1270 |
|  | R | 2163 |
| PAR | L | 502 |
|  | R | 552 |
| VIS | L | 1547 |
|  | R | 1600 |
| AUD | L | 486 |
|  | R | 442 |

### fMRI data acquisition and preprocessing

MRI scanning was performed on a 3-Tesla Siemens Skyra scanner with a 32-channel phased-array head coil at the imaging center at University of California, Davis. Functional data were collected using a T2-weighted echoplanar imaging (EPI) sequence. Each volume contained 60 axial slices (2.2mm thickness) parallel to the AC-PC line (TR, 1805ms; TE, 28ms). Each scan acquired 215 volumes (388s) and consisted of 40 visual search trials (fixed intertrial interval 2000ms) and 56 probe trials (jittered interstimulus interval with a mean of 4000ms). A total of 8 scans were acquired. A MPRAGE T1-weighted structural image (TR, 1800ms; TE, 2.97ms; 1✕1✕1mm^3^ resolution; 208 slices) was acquired for visualizing the associated anatomy. fMRI data were analyzed using SPM12 (Wellcome Trust Centre for Neuroimaging). The structural image was coregistered to the mean of the EPI images. Data preprocessing included slice timing correction and spatial realignment. We fitted the time series of each voxel with a general linear model (GLM) with 8 regressors (1 visual search condition and 7 probe conditions). The GLM also included regressors for each scan and 6 motion parameters.

### Representational similarity analysis

#### Behavioral representation of target templates

The primary question of interest was whether we could decode a biased target template that reflects individual decision biases in target identification based on visual search experiences. We hypothesized that this effect would be clearest in lateral prefrontal cortex given its adaptive coding of task relevance, but also tested parietal and visual regions known to be contain target representations (Bettencourt & Xu, 2016; Ester et al., 2009; Harrison & Tong, 2009). To do this, we performed representational similarity analysis (RSA) by extracting behavioral and neural representational geometries from each participant (Kriegeskorte et al., 2008; Nili et al., 2014). The behavioral target representation was measured by constructing a representational dissimilarity matrix (RDM) from the proportion of “target yes” responses in the probe trials (Figure 2A). The target-yes responses were false alarms when the probe color was a non-target color, and a hit when the color was a target color. The value in each cell of the RDM indicates the dissimilarity of the dependent measure between each pair of probe stimuli (Figure 2B). For example, if the proportion of target-yes responses to the -5° probe color was about the same as the 0° probe color, similarity would be high irrespective of the actual proportion of target-yes responses to the -5° and 0° probe color (Figure S2). We compared the brain RDM estimated from the patterns of neural activity in a *priori* ROIs with two candidate model RDMs. The model RDMs included (1) a biased RDM (*B*) constructed from each participants’ own probe performance (Figure 2B); (2) a veridical RDM (*V*) (Figure 2E), which assumed the target template was normally distributed and centered at the target color 0° (Figure 2D; *σ* = 7 and *μ* = 0) (Yu & Geng, 2019; Experiment 1; bidirectional group).

Because the two model RDMs were significantly correlated with each other (*M* ± 95% CI = .36 ± .10, z_19_ = 3.92, *p* < .0001), we tested for an effect of *B*(*V*) while controlling for the shared covariance between *B* and *V*, using partial correlation (Park et al., 2020). Specifically, we measured the extent to which the brain RDM estimated in each ROI was explained by *B’*(*V’*). For example, *B’* indicates the Euclidean distances between each pair of stimuli in *B* while regressing out its correlation with the pairwise Euclidean distances in *V*: *B’* = *B –* corr(*B, V*)\**V* (Figure 2C). After regressing out the partial correlation, the model RDMs were independent from each other while preserving high correlation with their original matrices (i.e., *B’* does not correlate with *V* while it still highly correlates with *B*).

#### Neural representation of target templates

A 7✕7 brain RDM for each ROI was constructed using the Euclidean distance based on the patterns of neural activity for each of the seven probe stimuli. We used *t*-statistics instead of regression estimates to down-weight response estimates from noisier voxels (Misaki et al., 2010; Walther et al., 2016). These *t* values were then normalized (mean response of each voxel across conditions = 0) to remove univariate differences between conditions. To ensure that the selected ROIs contained meaningful information regarding the template rather than rank noise, we performed a split half cross validation procedure. Specifically, we divided the imaging data into two independent splits of odd and even runs (four runs per split) and computed the representational distance between each pair of probe stimuli across the two splits. For example, the cross-validated distance between -5° and 0° was estimated as the Euclidean distance between the neural activity of -5° in one split and 0° in the other split. Because noise is independent between the two splits, the expected value of this distance is 0 if there is no systematic difference between the two conditions (Walther et al., 2016).

Next, we confirmed that the brain RDM of each ROI discriminated different probe stimuli with good sensitivity using the exemplar discriminability index (EDI) (Nili et al., 2020). EDI is defined as the mean of representational distances between different probe stimuli compared to the mean of the representational distances between the same probe stimuli (i.e., the average of the off diagonal vs. the average of the main diagonal). We confirmed that the EDI in all ROIs was positive (one sample *t* test, *ts* > 3, *ps* < .005), except for the control regions (*ps* > .10), suggesting that the different probe stimuli were discriminable in ROIs based on multivariate neural activity patterns.

The extent to which the brain RDM in each ROI was explained by the model RDMs was estimated with the Kendall’s τ_A_ rank correlation. The computed Kendall’s τ_A_ correlations were then subtracted from each participant’s baseline value, which was estimated as the mean of permuted rank correlations by randomizing stimulus labels 5000 times. For the group level inference, the significance of correlations was tested with the non-parametric Wilcoxon signed-rank test across participants (Nili et al., 2014). The resulting *p* values were corrected for multiple comparisons across the numbers of bilateral ROIs (*N* = 8), as well as the numbers of model RDMs (*N* = 2). We reported results corrected for family-wise error (FWE) with the Holm-Bonferroni method at *p* < 0.05, but stronger effects were indicated with asterisks.

#### Searchlight-based RSA

In order to be as inclusive as possible in searching for regions of the brain that might encode the target representations overall, we conducted a searchlight analysis (radius of 8mm) within the whole brain using the RSA toolbox (Nili et al., 2014) and custom MATLAB code. We correlated the brain RDM at each searchlight location with each participant’s model RDMs (*B* and *V*) using Kendall’s τ_A_ rank correlation. The results formed a continuous statistical map that show how well the model RDMs explained the brain RDM at the local brain regions. These images were further normalized, smoothed using an 8-mm full-width at half maximum (FWHM) Gaussian kernel and Fisher’s z transformed to confirm the statistical assumption (normality) required for the second-level parametric tests. Each participant’s statistical map was then submitted to a second-level one sample *t* test to find the voxels with correlation values greater than 0. The threshold for the resulting statistical maps were set at the cluster level *p* < .05, FWE corrected for multiple comparisons and a minimum number of voxels > 50.

### Drift diffusion model

The main analysis of the online experiment consisted of modeling visual search performance from the critical trials and probe performance (error rates and RT; Figure 5B and 5D) using a drift diffusion model. The separation between the two decision boundaries (*a*) and the non-decision time (*t*) were estimated as fitted free parameters that were the same across color distractor values for each participant while the drift rate (*v*) was estimated as a free parameter per color condition. Here, we were interested in how the drift rate (*v*), which characterizes the accumulation of noisy evidence over time until one of two decision boundaries is reached, differed across conditions (Ratcliff & McKoon, 2008). Drift rates in the critical search trials represent how easily the target can be distinguished from the distractor. Drift rates in the probe trials indicate how easily the probe stimulus can be identified as the target from memory. Higher drift rates indicate stronger evidence, whereas lower drift rates suggest weaker evidence.

All parameters were estimated using a hierarchical Bayesian parameter estimation method (Ratcliff & Childers, 2015). To perform hierarchical DDM, we used the Python-based toolbox, HDDM (Wiecki et al., 2013). The HDDM model was fit to accuracy-coded data (i.e., the upper and lower boundaries correspond to correct and incorrect responses, and the starting point was fixed at 0.5). For each participant’s data, we used Markov chain Monte Carlo (MCMC) sampling methods to estimate the posterior distribution of each parameter. Five chains were run with 10000 samples for each chain. The first 5000 warm-up samples were discarded as burn-in. Convergence was assessed by computing the Gelman-Rubin Ȓ statistic for each parameter. The range of R^^^ values across all group parameter estimates was between 0.99-1.10, suggesting that the samples of the different chains converged. Goodness of fit was visually inspected with the posterior predictive check method (Figure 5) (Wiecki et al., 2013).

## Acknowledgements

We would like to thank Phillip Witkowski for assistance in data collection, and Seongmin A. Park for helpful comments. This work was supported by NIH R01MH113855 to JJG. All data are publicly available in National Institute of Mental Health Data Archive (https://nda.nih.gov/edit_collection.html?id=2922).

## References

Badre, D., & Nee, D. E. (2018). Frontal Cortex and the Hierarchical Control of Behavior. Trends in Cognitive Sciences, 22(2), 170—188. https://doi.org/10.1016/j.tics.2017.11.005

Bae, G.-Y., Olkkonen, M., Allred, S. R., & Flombaum, J. I. (2015). Why some colors appear more memorable than others: A model combining categories and particulars in color working memory. Journal of Experimental Psychology. General, 144(4), 744—763. https://doi.org/10.1037/xge0000076

Baldauf, D., & Desimone, R. (2014). Neural Mechanisms of Object-Based Attention. 344, 5.

Bauer, B., Jolicoeur, P., & Cowan, W. B. (1996). Visual search for colour targets that are or are not linearly separable from distractors. Vision Research, 36(10), 1439—1466. https://doi.org/10.1016/0042-6989(95)00207-3

Becker, S. I. (2010). The Role of Target–Distractor Relationships in Guiding Attention and the Eyes in Visual Search. Journal of Experimental Psychology: General, 139(2), 247–265. https://doi.org/10.1037/a0018808

Bettencourt, K. C., & Xu, Y. (2016). Decoding the content of visual short-term memory under distraction in occipital and parietal areas. Nature Neuroscience, 19(1), 150—157. https://doi.org/10.1038/nn.4174

Bichot, N. P., Heard, M. T., DeGennaro, E. M., & Desimone, R. (2015). A Source for Feature-Based Attention in the Prefrontal Cortex. Neuron, 88(4), 832—844. https://doi.org/10.1016/j.neuron.2015.10.001

Bichot, N. P., Xu, R., Ghadooshahy, A., Williams, M. L., & Desimone, R. (2019). The role of prefrontal cortex in the control of feature attention in area V4. Nature Communications, 10(1). https://doi.org/10.1038/s41467-019-13761-7

Birman, D., & Gardner, J. L. (2019). A flexible readout mechanism of human sensory representations. Nature Communications, 10(1). https://doi.org/10.1038/s41467-019-11448-7

Bracci, S., Daniels, N., & Beeck H. O. de. (2017). Task Context Overrules Object-and Category-Related Representational Content in the Human Parietal Cortex. Cerebral Cortex (New York, NY), 27(1), 310—321. https://doi.org/10.1093/cercor/bhw419

Brainard, D. H. (1997). The Psychophysics Toolbox. Spatial Vision, 10(4), 433—436. https://doi.org/10.1163/156856897x00357

Bravo, M. J., & Farid, H. (2016). Observers change their target template based on expected context. Attention, Perception, \textbackslash& Psychophysics, 78(3), 829—837. https://doi.org/10.3758/s13414-015-1051-x

Bundesen, C. (1990). A theory of visual attention. Psychological Review, 97(4), 523—547. https://doi.org/10.1037/0033-295x.97.4.523

Castelhano, M. S., Pollatsek, A., & Cave, K. R. (2008). Typicality aids search for an unspecified target, but only in identification and not in attentional guidance. Psychonomic Bulletin & Review, 15(4), 795–801. https://doi.org/10.3758/PBR.15.4.795

Chelazzi, L., Duncan, J., Miller, E. K., & Desimone, R. (1998). Responses of Neurons in Inferior Temporal Cortex During Memory-Guided Visual Search. Journal of Neurophysiology, 80(6), 2918—2940. https://doi.org/10.1152/jn.1998.80.6.2918

Christophel, T. B., Klink, P. C., Spitzer, B., Roelfsema, P. R., & Haynes, J.-D. (2017). The Distributed Nature of Working Memory. Trends in Cognitive Sciences, 21(2), 111–124. https://doi.org/10.1016/j.tics.2016.12.007

Cunningham, C. A., & Wolfe, J. M. (2014). The role of object categories in hybrid visual and memory search. Journal of Experimental Psychology. General, 143(4), 1585—1599. https://doi.org/10.1037/a0036313

de la Vega, A., Yarkoni, T., Wager, T. D., & Banich, M. T. (2018). Large-scale Meta-analysis Suggests Low Regional Modularity in Lateral Frontal Cortex. Cerebral Cortex, 28(10), 3414–3428. https://doi.org/10.1093/cercor/bhx204

Desimone, R., & Duncan, J. (1995). Neural Mechanisms of Selective Visual Attention. Annual Review of Neuroscience, 18(1), 193—222. https://doi.org/10.1146/annurev.ne.18.030195.001205

D’Esposito, M., & Postle, B. R. (2015). The Cognitive Neuroscience of Working Memory. Annual Review of Psychology, 66(1), 115—142. https://doi.org/10.1146/annurev-psych-010814-015031

Duncan, J. (2001). An adaptive coding model of neural function in prefrontal cortex. Nature Reviews Neuroscience, 2(11), 820–829. https://doi.org/10.1038/35097575

Duncan, J. (2013). The Structure of Cognition: Attentional Episodes in Mind and Brain. Neuron, 80(1), 35–50. https://doi.org/10.1016/j.neuron.2013.09.015

Duncan, J., & Humphreys, G. W. (1989). Visual Search and Stimulus Similarity. 26.

Ester, E. F., Serences, J. T., & Awh, E. (2009). Spatially Global Representations in Human Primary Visual Cortex during Working Memory Maintenance. Journal of Neuroscience, 29(48), 15258—15265. https://doi.org/10.1523/jneurosci.4388-09.2009

Ester, E. F., Sprague, T. C., & Serences, J. T. (2015). Parietal and Frontal Cortex Encode Stimulus-Specific Mnemonic Representations during Visual Working Memory. Neuron, 87(4), 893—905. https://doi.org/10.1016/j.neuron.2015.07.013

Ester, E. F., Sutterer, D. W., Serences, J. T., & Awh, E. (2016). Feature-Selective Attentional Modulations in Human Frontoparietal Cortex. Journal of Neuroscience, 36(31), 8188— 8199. https://doi.org/10.1523/jneurosci.3935-15.2016

Feredoes, E., Heinen, K., Weiskopf, N., Ruff, C., & Driver, J. (2011). Causal evidence for frontal involvement in memory target maintenance by posterior brain areas during distracter interference of visual working memory. Proceedings of the National Academy of Sciences, 108(42), 17510—17515. https://doi.org/10.1073/pnas.1106439108

Funahashi, S., Bruce, C. J., & Goldman-Rakic, P. S. (1989). Mnemonic coding of visual space in the monkey’s dorsolateral prefrontal cortex. Journal of Neurophysiology, 61(2), 331— 349. https://doi.org/10.1152/jn.1989.61.2.331

Fuster, J. M., & Alexander, G. E. (1971). Neuron Activity Related to Short-Term Memory. Science, 173(3997), 652—654. https://doi.org/10.1126/science.173.3997.652

Geng, J. J., DiQuattro, N. E., & Helm, J. (2017). Distractor Probability Changes the Shape of the Attentional Template. Journal of Experimental Psychology: Human Perception and Performance, 43(12), 1993–2007. https://doi.org/10.1037/xhp0000430

Geng, J. J., & Witkowski, P. (2019). Template-to-distractor distinctiveness regulates visual search efficiency. Current Opinion in Psychology, 29, 119—125. https://doi.org/10.1016/j.copsyc.2019.01.003

Harrison, S. A., & Tong, F. (2009). Decoding reveals the contents of visual working memory in early visual areas. Nature, 458(7238), 632—635. https://doi.org/10.1038/nature07832

Hodsoll, J., & Humphreys, G. W. (2001). Driving attention with the top down: The relative contribution of target templates to the linear separability effect in the size dimension. Perception \textbackslash&#x026; Psychophysics, 63(5), 918—926. https://doi.org/10.3758/bf03194447

Hout, M. C., & Goldinger, S. D. (2015). Target templates: The precision of mental representations affects attentional guidance and decision-making in visual search. Attention, Perception \textbackslash& Psychophysics, 77(1), 128—149. https://doi.org/10.3758/s13414-014-0764-6

Kriegeskorte, N., Mur, M., & Bandettini, P. (2008). Representational Similarity Analysis – Connecting the Branches of Systems Neuroscience. Frontiers in Systems Neuroscience, 2. https://doi.org/10.3389/neuro.06.004.2008

Lee, S.-H., Kravitz, D. J., & Baker, C. I. (2013). Goal-dependent dissociation of visual and prefrontal cortices during working memory. Nature Neuroscience, 16(8), 997—999. https://doi.org/10.1038/nn.3452

Long, N. M., & Kuhl, B. A. (2018). Bottom-Up and Top-Down Factors Differentially Influence Stimulus Representations Across Large-Scale Attentional Networks. The Journal of Neuroscience, 38(10), 2495—2504. https://doi.org/10.1523/jneurosci.2724-17.2018

Malcolm, G. L., & Henderson, J. M. (2009). The effects of target template specificity on visual search in real-world scenes: Evidence from eye movements. Journal of Vision, 9(11), 8–8. https://doi.org/10.1167/9.11.8

Malcolm, G. L., & Henderson, J. M. (2010). Combining top-down processes to guide eye movements during real-world scene search. Journal of Vision, 10(2), 4–4. https://doi.org/10.1167/10.2.4

Miller, B. T., & D’Esposito, M. (2005). Searching for “the Top” in Top-Down Control. Neuron, 48(4), 535—538. https://doi.org/10.1016/j.neuron.2005.11.002

Miller, E. K., Erickson, C. A., & Desimone, R. (1996). Neural Mechanisms of Visual Working Memory in Prefrontal Cortex of the Macaque. Journal of Neuroscience, 16(16), 5154— 5167. https://doi.org/10.1523/jneurosci.16-16-05154.1996

Misaki, M., Kim, Y., Bandettini, P. A., & Kriegeskorte, N. (2010). Comparison of multivariate classifiers and response normalizations for pattern-information fMRI. NeuroImage, 53(1), 103—118. https://doi.org/10.1016/j.neuroimage.2010.05.051

Moore, T., Armstrong, K. M., & Fallah, M. (2003). Visuomotor Origins of Covert Spatial Attention. Neuron, 40(4), 671–683. https://doi.org/10.1016/S0896-6273(03)00716-5

Moore, T., & Zirnsak, M. (2017). Neural Mechanisms of Selective Visual Attention. Annual Review of Psychology, 68(1), 47—72. https://doi.org/10.1146/annurev-psych-122414-033400

Navalpakkam, V., & Itti, L. (2007). Search Goal Tunes Visual Features Optimally. Neuron, 53(4), 605—617. https://doi.org/10.1016/j.neuron.2007.01.018

Nili, H., Walther, A., Alink, A., & Kriegeskorte, N. (2020). Inferring exemplar discriminability in brain representations. PLOS ONE, 15(6), e0232551. https://doi.org/10.1371/journal.pone.0232551

Nili, H., Wingfield, C., Walther, A., Su, L., Marslen-Wilson, W., & Kriegeskorte, N. (2014). A Toolbox for Representational Similarity Analysis. PLoS Computational Biology, 10(4), e1003553. https://doi.org/10.1371/journal.pcbi.1003553

Panichello, M. F., & Buschman, T. J. (2021). Shared mechanisms underlie the control of working memory and attention. Nature, 592(7855), 601–605. https://doi.org/10.1038/s41586-021-03390-w

Park, S. A., Miller, D. S., Nili, H., Ranganath, C., & Boorman, E. D. (2020). Map Making: Constructing, Combining, and Inferring on Abstract Cognitive Maps. Neuron, 107(6), 1226-1238.e8. https://doi.org/10.1016/j.neuron.2020.06.030

Pelli, D. G. (1997). The VideoToolbox software for visual psychophysics: Transforming numbers into movies. Spatial Vision, 10(4), 437—442. https://doi.org/10.1163/156856897x00366

Peltier, C., & Becker, M. W. (2016). Decision processes in visual search as a function of target prevalence. Journal of Experimental Psychology: Human Perception and Performance, 42(9), 1466–1476. https://doi.org/10.1037/xhp0000248

Rainer, G., Asaad, W. F., & Miller, E. K. (1998). Selective representation of relevant information by neurons in the primate prefrontal cortex. Nature, 393(6685), 577–579. https://doi.org/10.1038/31235

Rajsic, J., & Woodman, G. F. (2020). Do we remember templates better so that we can reject distractors betterã Attention, Perception, & Psychophysics, 82(1), 269–279. https://doi.org/10.3758/s13414-019-01721-8

Ratcliff, R., & Childers, R. (2015). Individual Differences and Fitting Methods for the Two-Choice Diffusion Model of Decision Making. Decision (Washington, D.C.), 2015.

Ratcliff, R., & McKoon, G. (2008). The Diffusion Decision Model: Theory and Data for Two-Choice Decision Tasks. Neural Computation, 20(4), 873—922. https://doi.org/10.1162/neco.2008.12-06-420

Reynolds, J. H., & Heeger, D. J. (2009). The Normalization Model of Attention. Neuron, 61(2), 168—185. https://doi.org/10.1016/j.neuron.2009.01.002

Riggall, A. C., & Postle, B. R. (2012). The Relationship between Working Memory Storage and Elevated Activity as Measured with Functional Magnetic Resonance Imaging. The Journal of Neuroscience, 32(38), 12990—12998. https://doi.org/10.1523/jneurosci.1892-12.2012

Sarma, A., Masse, N. Y., Wang, X.-J., & Freedman, D. J. (2016). Task Specific versus Generalized Mnemonic Representations in Parietal and Prefrontal Cortices. Nature Neuroscience, 19(1), 143—149. https://doi.org/10.1038/nn.4168

Schaefer, A., Kong, R., Gordon, E. M., Laumann, T. O., Zuo, X.-N., Holmes, A. J., Eickhoff, S. B., & Yeo, B. T. T. (2018). Local-Global Parcellation of the Human Cerebral Cortex from Intrinsic Functional Connectivity MRI. Cerebral Cortex, 28(9), 3095—3114. https://doi.org/10.1093/cercor/bhx179

Scolari, M., Byers, A., & Serences, J. T. (2012). Optimal Deployment of Attentional Gain during Fine Discriminations. The Journal of Neuroscience, 32(22), 7723—7733. https://doi.org/10.1523/jneurosci.5558-11.2012

Serences, J. T., Ester, E. F., Vogel, E. K., & Awh, E. (2009). Stimulus-Specific Delay Activity in Human Primary Visual Cortex. Psychological Science, 20(2), 207—214. https://doi.org/10.1111/j.1467-9280.2009.02276.x

Squire, R. F., Noudoost, B., Schafer, R. J., & Moore, T. (2013). Prefrontal Contributions to Visual Selective Attention. Annual Review of Neuroscience, 36(1), 451–466. https://doi.org/10.1146/annurev-neuro-062111-150439

Sreenivasan, K. K., Vytlacil, J., & D’Esposito, M. (2014). Distributed and Dynamic Storage of Working Memory Stimulus Information in Extrastriate Cortex. Journal of Cognitive Neuroscience, 26(5), 1141—1153. https://doi.org/10.1162/jocn_a_00556

Swaminathan, S. K., & Freedman, D. J. (2012). Preferential encoding of visual categories in parietal cortex compared with prefrontal cortex. Nature Neuroscience, 15(2), 315–320. https://doi.org/10.1038/nn.3016

Treisman, A. M., & Gelade, G. (1980). A feature-integration theory of attention. Cognitive Psychology, 12(1), 97—136. https://doi.org/10.1016/0010-0285(80)90005-5

Walther, A., Nili, H., Ejaz, N., Alink, A., Kriegeskorte, N., & Diedrichsen, J. (2016). Reliability of dissimilarity measures for multi-voxel pattern analysis. NeuroImage, 137, 188—200. https://doi.org/10.1016/j.neuroimage.2015.12.012

Wiecki, T. V., Sofer, I., & Frank, M. J. (2013). HDDM: Hierarchical Bayesian estimation of the Drift-Diffusion Model in Python. Frontiers in Neuroinformatics, 7. https://doi.org/10.3389/fninf.2013.00014

Wolfe, J. M. (2021). Guided Search 6.0: An updated model of visual search. Psychonomic Bulletin & Review. https://doi.org/10.3758/s13423-020-01859-9

Yeo, B. T. T., Krienen, F. M., Sepulcre, J., Sabuncu, M. R., Lashkari, D., Hollinshead, M., Roffman, J. L., Smoller, J. W., Zöllei, L., Polimeni, J. R., Fischl, B., Liu, H., & Buckner, R. L. (2011). The organization of the human cerebral cortex estimated by intrinsic functional connectivity. Journal of Neurophysiology, 106(3), 1125—1165. https://doi.org/10.1152/jn.00338.2011

Yu, X., & Geng, J. J. (2019). The Attentional Template Is Shifted and Asymmetrically Sharpened by Distractor Context. Journal of Experimental Psychology: Human Perception and Performance, 45(3), 336–353. https://doi.org/10.1037/xhp0000609

