## Supplemental Materials for "Pattern similarity in the frontoparietal control network reflects an “off-veridical” template that optimizes target-match decisions during visual search"

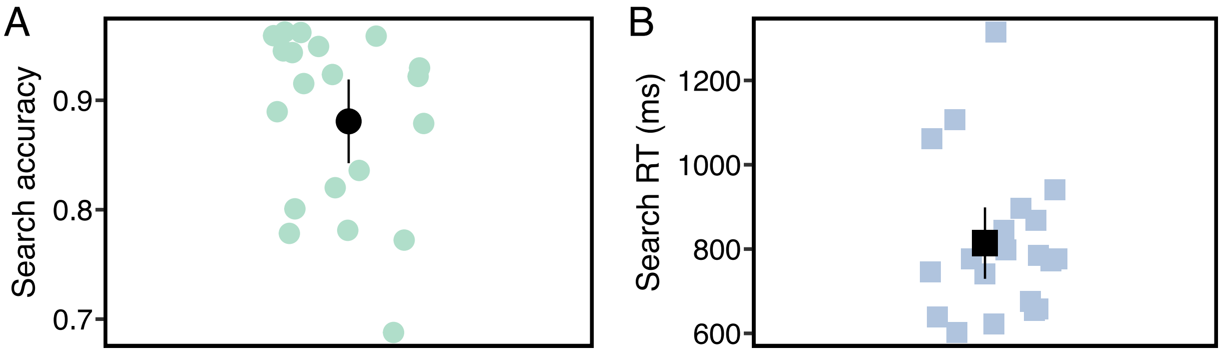


*Figure S1*. Visual search accuracy and RTs from the visual search trials. The colored dots represent individual data points, and the black ones indicate the mean values. All error bars are the 95% confidence intervals.

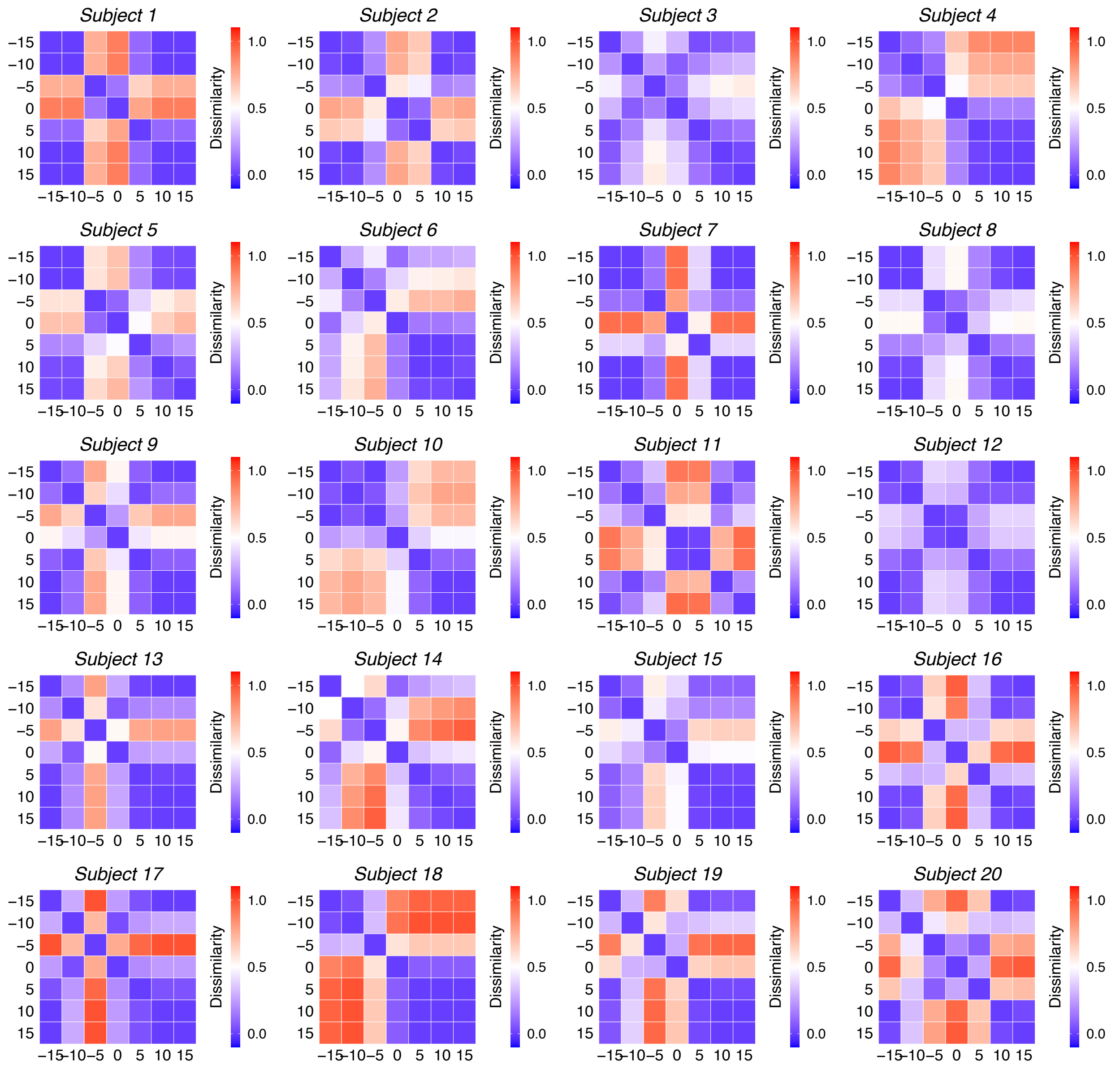


*Figure S2*. The individual biased behavioral RDM (*B*) generated by participants' own probe task performance.

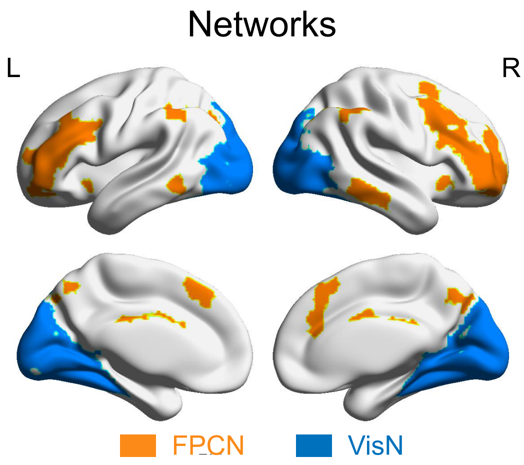


*Figure S3*. Two independently defined resting-state networks of a priori interest: frontoparietal control network (FPCN) and visual networks (VisN) (Yeo et al., 2011).

**Button response RDM searchlight**

The significant searchlight results observed in regions related to motor and somatosensory functions in the left hemisphere (contralateral to the right index and middle fingers that were used for button presses) raised the possibility that the selected ROIs were driven in part by the motor plan of making an index or middle finger button response, which corresponded to the yes/no target-match decision. Although the full $B'$ and $V'$ matrices cannot be explained by motor movements alone, to address this, we created a dissimilarity matrix that modeled the button responses *per se*, regardless of the actual probe stimulus. Specifically, each trial was labeled as either a "yes" trial or "no" trial based on the individual's response. The group of "yes" and "no" trials were randomly split into two halves to create a response model RDM in which trials with the same response, irrespective of the actual stimulus, were maximally similar whereas trials with different responses were maximally dissimilar (Figure S4): the dissimilarity distance between the same responses was 0, and the distance between the different response types was 1. We then performed another whole brain searchlight for each participant using their individual button response RDM as the reference model. The group analysis identified only one significant region ($p < .001$, uncorrected) located in left PoG (Figure S4; [x, y, z] = [-44, -18, 58], $z_{19} = 3.53$, # of voxels = 92) in motor cortex contralateral to the right hand. 39 and 37 number of voxels in this cluster were overlapped with left PrG identified from the searchlight map with the biased and veridical RDM, respectively. The results support the notion that motor regions, e.g., left PrG, captured by the two RDMs of interest were involved in the actual motor response rather than response independent decision processes.

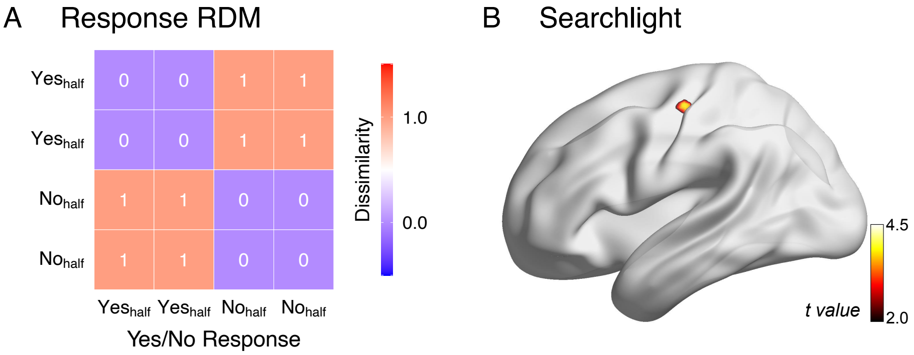


*Figure S4*. A) The button response behavioral RDM. B) Brain maps showing the clusters identified in the whole brain searchlight analysis, located in the left PoG.
